# “Comparison of antibiotic protein binding in human plasma vs. rabbit plasma”

**DOI:** 10.1101/2023.03.23.534048

**Authors:** Maximilian Pesta, Philip Datler, Georg Scheriau, Peter Wohlrab, Sabine Eberl, Edith Lackner, Claudia Franz, Walter Jäger, Alexandra Maier-Salamon, Markus Zeitlinger, Edda Tschernko

## Abstract

Rabbits are frequently used for the examination of the pharmacokinetics and effectiveness of antibiotic substances. However, antibiotics vary substantially in protein binding affecting the concentration of the antimicrobially effective unbound drug. We hypothesized that the binding properties of vancomycin, meropenem and ceftriaxone might vary between human and rabbit plasma. In an in-vitro study we observed dose dependent variability in protein binding of antibiotics between species. Thus, in-vitro-pre-studies are required to guarantee for translational conditions.

## Introduction

Small animals are frequently used for translational studies of antibiotic efficacy. Numerous antibiotics show high protein binding, mainly to albumin, but only the unbound substance is antimicrobially active. In addition, tissue distribution and renal clearance might be effected by protein binding. Therapeutically effective unbound plasma concentrations of vancomycin, meropenem and ceftriaxone are influenced by several factors (1,2). To date, few studies are available comparing antibiotic binding of human and rabbit albumin for various antibiotic substances (3, 4). However, none of the studies evaluated protein binding in human and in rabbit plasma comparing normal antibiotic concentrations with extremely high or low antibiotic concentrations. Differences in protein binding properties at variable antibiotic concentrations could define the antibiotic concentration range in which translational comparability between species can be expected. Therefore, we undertook an in-vitro study comparing the binding properties of three antibiotics with different albumin binding affinity. We hypothesized that the binding properties of vancomycin, meropenem and ceftriaxone might vary between human and rabbit plasma. In an in-vitro study, we set out to identify dose-dependent variability in protein binding.

Microdialysis technique (5) was used for determination of protein binding of vancomycin, meropenem and ceftriaxone, since these antibiotics are known to have different binding properties to albumin in humans. Three concentrations were used for each antibiotic substance: expected peak antibiotic plasma concentration after standard treatment as quoted in the literature, 25% and 1000% of antibiotic standard concentration of established treatment concentration. For vancomycin we used concentrations of 5 µg/ml, 20 µg/ml and 200 µg/ml, (6) for meropenem we used concentrations of 2.5 µg/ml, 10 µg/ml and 100 µg/ml (7), and for ceftriaxone we used concentrations of 4 µg/ml, 15 µg/ml and 150 µg/ml (8). Each drug concentration was emersed in three solutions: NaCl 0.9%, human citrat plasma and rabbit citrat plasma. The concentration of vancomycin, meropenem and ceftriaxone in microdialysate samples was determined by HPLC using a Dionex “UltiMate 3000” system (Dionex Corp., Sunnyvale, CA) with UV detection at 260 nm, 296 nm and 220 nm, respectively. Chromatographic separation for all drugs was carried out on a Hypersil BDS-C18 column (5 µm, 250 × 4.6 mm I.D., Thermo Fisher Scientific, Inc, Waltham, MA), preceded by a Hypersil BDS-C18 precolumn (5 µm, 10 × 4.6mm I.D.), constantly heated to 45°C. Calibration of the chromatograms for all three drugs was accomplished using the external standard method by spiking drug-free microdialysate with standard solutions of the respective drugs. The limit of quantification for vancomycin, meropenem and ceftriaxone was 0.1 µg/ml; with coefficients of accuracy and precision below 8.7%.

## Results

Vancomycin protein binding in rabbit and human plasma were comparable in the low antibiotic vancomycin concentration of 5 µg/mL (39.0±16.0% vs 43.7±13.5; p=0.612; Figure) and standard treatment plasma concentration of 20 µg/mL (38.2±14.0% vs 39.8±13.2%; p=0.71; Figure). In contrast, in high plasma vancomycin concentrations of 200 µg/mL was significantly lower protein binding in rabbit plasma vs human plasma (43.1±10.1% vs 54.0±9.0%; p=0.002; Figure).

Ceftriaxone protein binding in rabbit and human plasma were 41.9±7.7% vs 45.4±10% in standard treatment plasma concentrations of 15 µg/mL (p=0.24; Figure) and 37.1±6.8% vs 39.8±7.4% in high plasma ceftriaxone concentrations of 150 µg/mL (p=0.27; Figure). In contrast, in low plasma ceftriaxone concentrations of 4 µg/mL (p<0.001; Figure) rabbit plasma exhibited significant lower protein binding vs human plasma (27.3±12.3% vs 40.3±9.8%; p<0.05; Figure).

Meropenem protein binding in rabbit and human plasma were 13.8±9.5% vs 17.3±10.2% in low antibiotic meropenem plasma concentration of 2.5 µg/mL (p=0.66; Figure) and in high antibiotic meropenem plasma concentration of 100 µg/mL 9.7±4.6% vs 11.1±5.6% (p=0.43; Figure). However, in the standard treatment plasma concentrations of 10 µg/mL significantly lower protein binding was displayed in rabbit plasma compared to human plasma (11.9±7.3% vs 26.6±11.6%; p<0.001; Figure). Meropenem measurements of protein binding were carried out twice in different laboratories to doublecheck and validate the primary findings. Results in both measurement series were found to be comparable.

## Conclusion

For vancomycin and ceftriaxone standard treatment plasma concentrations were comparable between species. Extreme plasma concentrations of these antibiotics revealed differences in antibiotic binding between species. Remarkably, meropenem protein binding was detected to be significantly different between rabbit and human plasma if standard antibiotic treatment concentrations are aimed for. Thus, rabbit models lack translational value for studies with meropenem in standard concentrations. Since protein binding can be substantially different at high, normal and low antibiotic concentrations in plasma depending on the antibiotic substance examined, it is advisable to perform in-vitro studies with the respective drug under investigation before animal studies are carried out. Animal species and drug concentrations must be chosen carefully before performing translational antibiotic research, because antibiotic protein binding can vary substantially between species and plasma concentrations. Thus, in-vitro pre-studies of antibiotic binding can be essential before choosing a species and an antibiotic substance at a certain concentration in order to guarantee for comparability to human conditions.

---

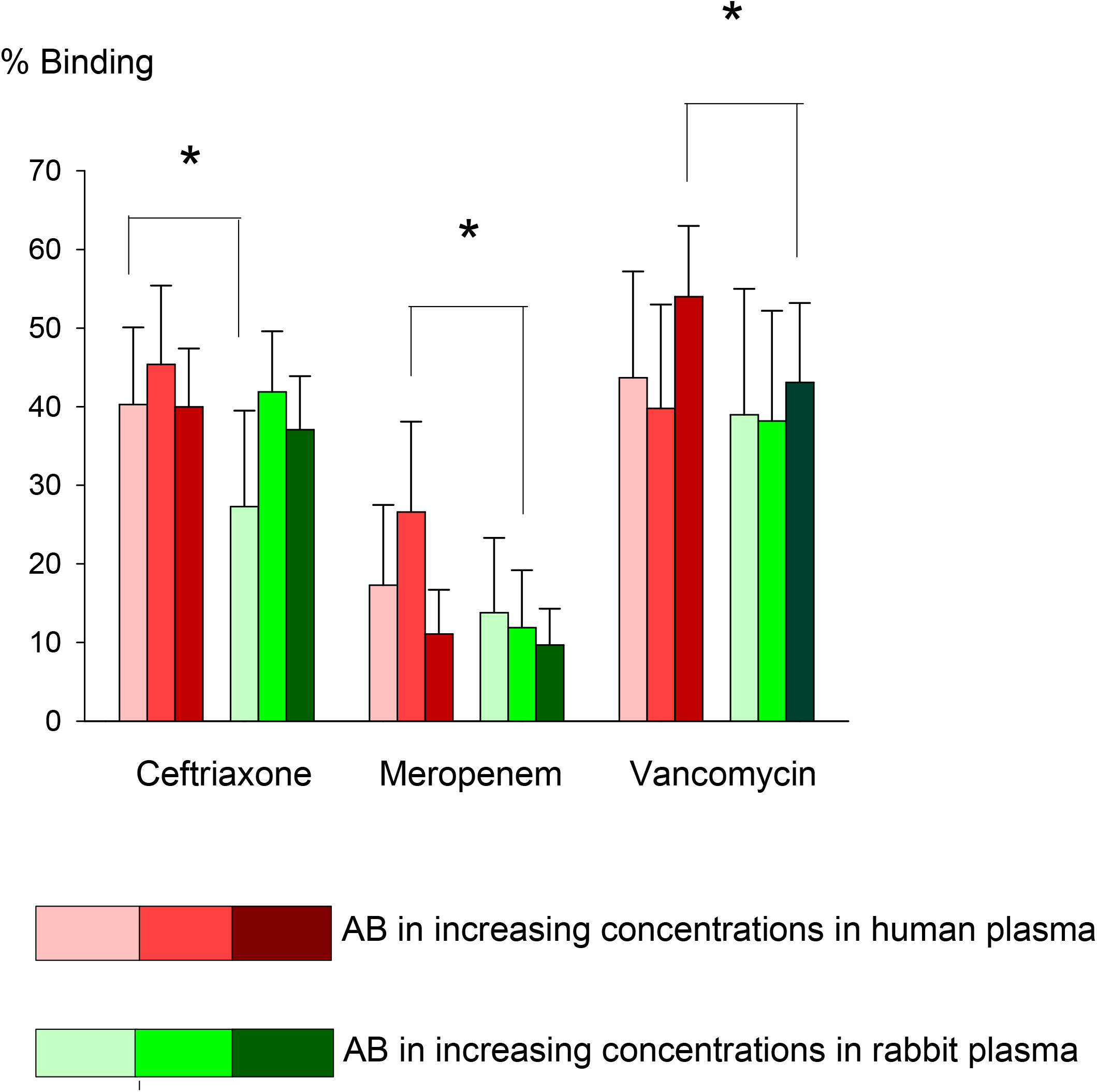
Shades of red indicate in-vitro human plasma experiments and shades of green indicate in-vitro rabbit plasma experiments. Examined antibiotic concentrations increase from left to right or from light to dark colors, respectively. P<0.05 is indicated with an asterisk.

